# DDX3 depletion represses translation of mRNAs with complex 5′ UTRs

**DOI:** 10.1101/589218

**Authors:** Lorenzo Calviello, Srivats Venkataramanan, Karol J. Rogowski, Emanuel Wyler, Kevin Wilkins, Malvika Tejura, Bao Thai, Jacek Krol, Witold Filipowicz, Markus Landthaler, Stephen N. Floor

**Affiliations:** Department of Cell and Tissue Biology, University of California, San Francisco, San Francisco, California, USA, 94143; Berlin Institute for Medical Systems Biology, Max-Delbrück-Center for Molecular Medicine in the Helmholtz Association, 13125, Berlin, Germany; Institute of Molecular and Clinical Ophthalmology Basel, Basel, Switzerland; Friedrich Miescher Institute for Biomedical Research, Basel, Switzerland; IRI Life Sciences, Institute für Biologie, Humboldt Universität zu Berlin, Philippstraße 13, 10115, Berlin, Germany; Helen Diller Family Comprehensive Cancer Center, University of California, San Francisco, San Francisco, California, USA, 94143

**Keywords:** translational control, DEAD-box proteins, RNA, PAR-CLIP, ribosome profiling, post-transcriptional control

## Abstract

DDX3 is an RNA chaperone of the DEAD-box family that regulates translation. Ded1, the yeast ortholog of DDX3, is a global regulator of translation, whereas DDX3 is thought to preferentially affect a subset of mRNAs. However, the set of mRNAs that are regulated by DDX3 are unknown, along with the relationship between DDX3 binding and activity. Here, we use ribosome profiling, RNA-seq, and PAR-CLIP to define the set of mRNAs that are regulated by DDX3 in human cells. We find that while DDX3 binds highly expressed mRNAs, depletion of DDX3 particularly affects the translation of a small subset of the transcriptome. We further find that DDX3 binds a site on helix 16 of the human ribosome, placing it immediately adjacent to the mRNA entry channel. Translation changes caused by depleting DDX3 levels or expressing an inactive point mutation are different, consistent with different association of these genetic variant types with disease. Taken together, this work defines the subset of the transcriptome that is responsive to DDX3 inhibition, with relevance for basic biology and disease states where DDX3 is altered.

## Introduction

Translation initiation is affected by mRNA regulatory elements. The DEAD-box RNA chaperone DDX3 and its yeast ortholog Ded1 facilitate translation initiation on mRNAs with structured 5′ untranslated regions (UTRs) (Guenther et al. 2018; Lai et al. 2008; Oh et al. 2016; Soto-Rifo et al. 2012), a function that is essential in all eukaryotes (Sharma and Jankowsky 2014). Dysfunction in DDX3 is linked to numerous diseases and cancers, including medulloblastoma (Epling et al. 2015; Floor et al. 2016; Jones et al. 2012; Kool et al. 2014; Oh et al. 2016; Pugh et al. 2012; Robinson et al. 2012; Valentin-Vega et al. 2016), many other cancer types (Sharma and Jankowsky 2014), and *de novo* developmental delay (Deciphering Developmental Disorders 2015; Snijders Blok et al. 2015; Wang et al. 2018; Lennox et al. 2020). Previous work studied how translation is altered by DDX3 variants found in medulloblastoma (Oh et al. 2016; Valentin-Vega et al. 2016), which are exclusively missense variants that preferentially target conserved residues. In contrast, hematological cancers like natural killer/T-cell lymphoma (Dufva et al. 2018; Jiang et al. 2015) and others (Wang et al. 2011; Schmitz et al. 2012) also have frequent variants in *DDX3X*, but they are mostly truncating or frameshift variants resulting in decreased expression. Changes in gene expression occurring as a result of decreased DDX3 levels remain incompletely understood.

Inactivation of Ded1 in yeast leads to polysome collapse and global downregulation of translation (Chuang et al. 1997; de la Cruz et al. 1997). More recent work showed that Ded1 is required for translation of most transcripts in yeast using genome-wide approaches (Guenther et al. 2018; Sen et al. 2015). In contrast, DDX3 depletion seems to only affect translation of a subset of expressed transcripts (Ku et al. 2018; Lai et al. 2008, 2010; Lee et al. 2008; Soto-Rifo et al. 2012). Despite the importance of DDX3 to normal function and its alteration in diverse disease states, the set of genes that depend on DDX3 for translation is not clearly defined. Moreover, it has been challenging to relate DDX3 binding to functional effects on bound mRNAs, and it was unclear if DDX3 is functioning outside of translation initiation given that binding was detected in coding sequences and 3′ UTRs (Oh et al. 2016; Valentin-Vega et al. 2016).

Here, we depleted DDX3 protein levels and measured alterations to translation and RNA abundance using ribosome profiling and RNA-seq. We also characterized DDX3 binding by PAR-CLIP, exploiting the presence of T>C mutations as a diagnostic hallmark of protein-RNA interactions. We observed robust interactions between DDX3 and transcript 5′ UTRs, as well as a specific and conserved site on the 40S ribosomal subunit. We found that transcripts with structured 5′ UTRs are preferentially affected by DDX3. We used an *in vitro* reporter system to conclude that decreases in ribosome occupancy upon DDX3 depletion are driven by 5′ UTRs. Taken together, our results support a model for DDX3 function where interactions with the small ribosomal subunit facilitate translation on messages with structured 5′ UTRs, which, when inactivated, pathologically deregulates protein synthesis.

## Methods

### NGS data pre-processing

Ribo-seq fastq files were stripped of the adapter sequences using cutadapt. UMI sequences were removed and reads were collapsed to fasta format. Reads were first aligned against rRNA (accession number U13369.1), and to a collection of snoRNAs, tRNAs and miRNA (retrieved using the UCSC table browser) using bowtie2 in the ‘local’ alignment mode.

Remaining reads were mapped to the hg38 version of the genome (without scaffolds) using STAR 2.6.0a (Dobin et al. 2013) supplied with the GENCODE 32 .gtf file. A maximum of 3 mismatches and mapping to a maximum of 50 positions was allowed. De-novo splice junction discovery was disabled for all datasets. Only the best alignment per each read was retained. Read counts for all libraries are in Table S3.

### PAR-CLIP peak calling

Peak calling for PAR-CLIP reads was performed with PARalyzer v1.5 (Corcoran et al. 2011) in the “EXTEND_BY_READ” mode using the following parameters: BANDWIDTH=3

CONVERSION=T>C

MINIMUM_READ_COUNT_PER_GROUP=5

MINIMUM_READ_COUNT_PER_CLUSTER=5

MINIMUM_READ_COUNT_FOR_KDE=5

MINIMUM_CLUSTER_SIZE=8

MINIMUM_CONVERSION_LOCATIONS_FOR_CLUSTE R=1

MINIMUM_CONVERSION_COUNT_FOR_CLUSTER=3

MINIMUM_READ_COUNT_FOR_CLUSTER_INCLUSIO N=5

MINIMUM_READ_LENGTH=18

MAXIMUM_NUMBER_OF_NON_CONVERSION_MISM ATCHES=0

Peaks with more than 10 reads were retained for subsequent analysis.

Coverage-normalized T>C conversions on rRNA for positions with 2000 reads or more (Figure S3A) were mapped onto the 18S rRNA sequence from PDB entry 6FEC and visualized using UCSF Chimera (Pettersen et al. 2004).

### Differential expression analysis

Count matrices for Ribo-seq and RNA-seq were built using reads mapping uniquely to CDS regions of protein-coding genes, using the Bioconductor packages *GenomicFeatures, GenomicFiles* and *GenomicAlignments*. Genomic and transcript regions where extracted using *Ribo-seQC* (Calviello et al. 2019). Only reads mapping for more than 25nt were used. Differential analysis was using DESeq2 (Love et al. 2014). Concordant changes were defined using an FDR cutoff of 0.01 for RNA-seq and Ribo-seq individually and ensuring the same directionality in the estimated fold changes.

Changes in translation efficiency were calculated using DESeq2 by using assay type (RNA-seq or Ribo-seq) as an additional covariate. Translationally regulated genes were defined using an FDR cutoff of 0.05 from a likelihood ratio test, using a reduced model without the assay type covariate, e.g. assuming no difference between RNA-seq and Ribo-seq counts (Chothani et al. 2019).

For both RNA-seq and Ribo-seq, only genes with BaseMean >8 or more than the bottom 10% of the library were used. GO enrichment analysis was performed with the *topGO* package, using the fisher test with default parameters.

The Random Forest regression was run using the *randomForest* package with default parameters, using the following features for each gene:

- TPM values using RNA-seq (in log scale);
- Baseline TE levels, defined as ratio of Ribo to RNA reads;
- Baseline RNA mature levels, defined as length-normalized ratio of RNA-seq reads in introns vs. exons;
- GC content, length (in log scale) and ribosome density in: 5′ UTRs, a window of 25nt around start and stop codons, CDS regions, non-coding internal exons, introns, and 3′ UTRs;
- Additional sequence features, including density of motif scores (calculated using the *Biostrings* package) for the following motifs, partially taken from (Murat et al. 2018):

- TOP: a binary variable indicating if gene belongs to core TOP mRNA, as defined in (Philippe et al. 2020)
- PRTE(pyrimidine-rich translational element): “[CU][CU][CU][CU][U][CU][CU][CU]”
- TISU(Translator Initiator of Short 5’-UTR): “[CG][A][A][CG][A][U][G][G][C][G][G][C]”
- CERT (cytosine-enriched regulator of translation): “[CG][CGU][CGU][CG][CGU][C][CGU][C][C A][GU][C][CGU][CGUA][CG][C]”
- PQS: propensity to create G quadruplexes, calculated using the *pqsfinder* R package (Hon et al. 2017)

Feature importance (measured by mean decrease in accuracy) and correlation between predicted and measured test data were calculated on a 10-fold cross-validation scheme.

### Meta-transcript profiles

PARalyzer peaks (and peaks from the POSTAR2 repository (Zhu et al. 2019)) were mapped on transcript coordinates using one coding transcript per gene: such transcript was chosen to have the longest 5′ UTR and the most common annotated start codon for that gene. Transcript positions were converted into bins using 15 bins for each 5′ UTR, 30 bins for each CDS and 20 bins for each 3′ UTR. Peak scores were normalized for each transcript (to sum up to 1), and values were summed for each bin to build aggregate profiles. When plotting profiles for different RBPs. the aggregate profiles were further normalized to a sum of 1. To build the average meta-transcript profile in Figure 4C, conversion specificity values were averaged per transcript bin. To create shuffled profiles in Figure 4 and S3, 5 random positions for each peak were taken from the same bound UTR.

### Additional transcript features analysis

To compare read mapping locations within transcripts, a window of 25nt around the start codon was subtracted from annotated 5′ UTRs and CDS. 5′ UTRs and CDS regions in genomic and transcriptomic space were retrieved using *Ribo-seQC*. Counts on 5′ UTR and CDS were first averaged between replicates. The ratio 5′ UTR to CDS of these counts were calculated for each gene, in the siRNA and controls condition. The log2 of the ratio siDDX3/control for those values represents the skew of counts towards 5′ UTR in the siDDX3 condition:

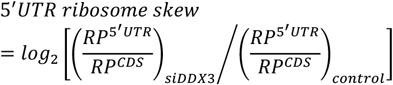

RNA *in silico* folding was performed on 5′ UTRs sequences using *RNAlfold* (Lorenz et al. 2016) with default parameters. Average ΔG values per nucleotide were calculated averaging the ΔG values of each structure overlapping that nucleotide. %GC content and T>C transition specificity (defined as *ConversionEventCount* / (*ConversionEventCount*+*NonConversionEventCount*)) for each PAR-CLIP peak were derived using the clusters.csv output file from PARalyzer. Gviz was used to plot tracks for RNA-seq, Ribo-seq and PAR-CLIP over different transcripts.

Source code to reproduce all the figures and can be found at: https://github.com/lcalviell/DDX3X_RPCLIP

### Ribosome profiling

Knockdown RP: Flp-In T-REx HEK293 cells transfected with control siPool and with *DDX3X*-targeting siPools (siTOOLs Biotech) were washed with PBS containing 100 μg/ mL cycloheximide, flash frozen on liquid nitrogen, lysed in lysis buffer (20 mM TRIS-HCl pH 7.4, 150mM NaCl, 5 mM MgCl2 1 % (v/v) Triton X-100, 25 U/ mL TurboDNase (Ambion), harvested, centrifuged at 20,000 g for 4 min at 4°C and supernatants were stored at −80°C. Thawed lysates were treated with RNase I (Ambion) at 2.5 U/μL for 45 min at room temperature with slow agitation. Further RNase activity was stopped by addition of SUPERase:In (Ambion). Next Illustra MicroSpin Columns S-400 HR (GE Healthcare) were used to enrich for ribosome complexes. RNA was extracted from column flow throughs with TRIzol (Ambion) reagent. Precipitated nucleic acids were further purified and concentrated with Zymo-Spin IIC column (Zymo Research). Obtained RNA was depleted of rRNAs with Ribo-Zero Gold Kit (Human/Mouse/Rat) kit (Illumina), separated in 17% Urea gel and stained with SYBR Gold (Invitrogen). Gel slices containing nucleic acids 27 to 30 nucleotides long were excised and incubated in a thermomixer with 0.3 M NaCl at 4°C overnight with constant agitation to elute RNA. After precipitation nucleic acids were treated with T4 polynucleotide kinase (Thermo Scientific). Purified RNA was ligated to 3′ and 5′ adapters, reverse transcribed and PCR amplified. The amplified cDNA was sequenced on a HiSeq2000 (Illumina).

Degron RP: *DDX3X*-mAID tagged HCT-116 (one 15 cm dish at 80-90% confluency per replicate) cells expressing OsTIR1 were transfected with either wild-type *DDX3X* or *DDX3X* R326H. 24 hours post-transfection, media was changed and fresh media with 500 μM indole 3-acetic acid (IAA) was added to cells. Un-transfected cells were treated with either DMSO or IAA. 48 hours after auxin addition, cells were treated with 100 μg/mL cycloheximide (CHX) and harvested and lysed as described in (McGlincy and Ingolia 2017). Briefly, cells were washed with PBS containing 100 μg/ mL CHX and lysed in ice-cold lysis buffer (20 mM TRIS-HCl pH 7.4, 150mM NaCl, 5 mM MgCl2, 1mM DTT, 100 μg/ mL CHX, 1 % (v/v) Triton X-100, 25 U/ mL TurboDNase (Ambion). 240 μl lysate was treated with 6 μl RNase I (Ambion, 100 U/μl) for 45 minutes at RT with gentle agitation and further digestion halted by addition of SUPERase:In (Ambion). Illustra Microspin Columns S-400 HR (GE healthcare) were used to enrich for monosomes, and RNA was extracted from the flow-through using Direct-zol kit (Zymo Research). Gel slices of nucleic acids between 24-32 nts long were excised from a 15% urea-PAGE gel. Eluted RNA was treated with T4 PNK and preadenylated linker was ligated to the 3′ end using T4 RNA Ligase 2 truncated KQ (NEB, M0373L). Linker-ligated footprints were reverse transcribed using Superscript III (Invitrogen) and gel-purified RT products circularized using CircLigase II (Lucigen, CL4115K). rRNA depletion was performed using biotinylated oligos as described in (Ingolia et al. 2012) and libraries constructed using a different reverse indexing primer for each sample.

### PAR-CLIP Experiments

Flp-In T-REx HEK293 cells expressing FLAG/HA-tagged *DDX3X* (Spitzer et al. 2013) were labelled with 100 μM 4-thiouridine for 16h. PAR-CLIP was performed generally as described (Hafner et al. 2010; Zarnegar et al. 2016). Briefly, cells were UV-crosslinked with 0.15 J/cm^2^ at 365nm, and stored at −80 °C. Obtained cell pellets were lysed in 3 times the cell pellet volume of NP-40 lysis buffer (50 mM HEPES-KOH at pH 7.4, 150 mM KCl, 2mM EDTA, 1mM NaF, 0.5% (v/v) NP-40, 0.5 mM DTT, complete EDTA-free protease inhibitor cocktail (Roche)), incubated 10 min on ice and centrifuged at 13,000 rpm for 15 min at 4 °C. Supernatants were filtered through 5 μm syringe filter. Next, lysates were treated with RNase I (Thermo-Fisher Scientific) at final concentration of 0.25U/μL for 10 min at room temperature. Immunoprecipitation of the DDX3/RNA complexes was performed with FLAG magnetic beads (Sigma). After IP and washing, the protein-bound RNAs were 3′ de-phosphorylated and 5′-end phosphorylated using T4 PNK with 0.01% Triton X-100, and the NIR fluorescent adaptor (5′-OH-AGATCGGAAGAGCGGTTCAGAAAAAAAAAAAA/iAzid eN/AAAAAAAAAAAA/3Bio/-3′) was ligated to the RNA using truncated RNA ligase 2 K227Q (NEB) overnight at 16°C, shaking at 1600 rpm. Crosslinked protein–RNA complexes were resolved on a 4–12% NuPAGE gel (Thermo-Fisher Scientific) and transferred to a nitrocellulose membrane. Protein–RNA complex migrating at an expected molecular weight were excised, and RNA by proteinase K (Roche) treatment and phenol–chloroform extraction. Purified RNA was further ligated to 5′ adapters, reverse transcribed and PCR amplified. The amplified cDNA was sequenced on a NextSeq 500 device (Illumina).

### *In vitro* transcription, capping, and 2′-O methylation of reporter RNAs

Annotated 5′ UTRs for selected transcripts were cloned upstream of Renilla Luciferase (RLuc) under the control of a T7 promoter, with 60 adenosine nucleotides downstream of the stop codon to mimic polyadenylation. 5′ UTR sequences are in Table S4. Untranslated regions were cloned using synthetic DNA (Integrated DNA Technologies) or by isolation using 5′ RACE (RLM-RACE kit, Invitrogen). Template was PCR amplified using Phusion polymerase from the plasmids using the following primers, and gel purified, as described (Floor and Doudna 2016).

pA60 txn rev: TTT TTT TTT TTT TTT TTT TTT TTT TTT TTT TTT TTT TTT TTT TTT TTT TTT TTT TTT TTT CTG CAG

pA60 txn fwd: CGG CCA GTG AAT TCG AGC TCT AAT ACG ACT CAC TAT AGG

100 μL *in vitro* transcription reactions were set up at room temperature with 1-5 micrograms of purified template, 7.5mM ACGU ribonucleotides, 30mM Tris-Cl pH 8.1, 125mM MgCl_2_, 0.01% Triton X-100, 2mM spermidine, 110mM DTT, T7 polymerase and 0.2 U/μL units of Superase-In RNase inhibitor (Thermo-Fisher Scientific). Transcription reactions were incubated in a PCR block at 37 °C for 1 hour. 1 μL of 1 mg/mL pyrophosphatase (Roche) was added to each reaction, and the reactions were subsequently incubated in a PCR block at 37 °C for 3 hours. 1 unit of RQ1 RNase-free DNase (Promega) was added to each reaction followed by further incubation for 30 minutes. RNA was precipitated by the addition of 200 μL 0.3M NaOAc pH 5.3, 15 μg GlycoBlue co-precipitant (Thermo-Fisher Scientific) and 750 μL 100% EtOH. Precipitated RNA was further purified over the RNA Clean & Concentrator-25 columns (Zymo Research). Glyoxal gel was run to assess the integrity of the RNA before subsequent capping and 2′ O-methylation.

20 μg of total RNA was used in a 40 μL capping reaction with 0.5mM GTP, 0.2mM s-adenosylmethionine (SAM), 20 units of Vaccinia capping enzyme (New England Biolabs), 100 units of 2′-O-Me-transferase (New England Biolabs) and 25 units RNasin Plus RNase inhibitor (Promega). The reactions were incubated at 37 °C for 1 hour, followed by purification over the RNA Clean & Concentrator-25 columns (Zymo Research) and elution in DEPC H_2_O. Glyoxal gel was run to assess the integrity of the RNA before proceeding to *in vitro* translation reactions.

### Transfection of siRNA for *in vitro* translation

HEK293T cells in 150mM plates were transfected with 20 μL of siRNA (against DDX3 or a non-targeting control) using Lipofectamine 2000 (Thermo Fisher Scientific), following manufacturer’s instructions. Cells were harvested for preparation of cellular extracts after 48 hours.

### Generation of DDX3 mutant translation extracts

DDX3 WT and R326H mutant constructs were synthesized and cloned downstream of a CMV promoter (Twist Biosciences). 40 μg of plasmids were transfected into HCT116 cells using Lipofectamine 2000 (Thermo Fisher Scientific), following manufacturer’s instructions. Cells were treated with 500μM Indole-3-acetic acid (IAA) 24 hours post-transfection and harvested for preparation of cellular extracts after a further 48 hours.

### Preparation of cellular extracts for *in vitro* translation

Three to five 150mm plates of HEK293T or HCT116 cells were trypsinized and pelleted at 1000g, 4 °C. One cell-pellet volume of lysis buffer (10mM HEPES, pH 7.5, 10mM KOAc, 0.5mM MgOAc_2_, 5mM DTT, and 1 tablet Complete mini EDTA free protease inhibitor (Sigma) per 10 mL) was added to the cell pellet and was incubated on ice for 45 minutes. The pellet was homogenized by trituration through a 26G needle attached to a 1 mL syringe 13-15 times. Efficiency of disruption was checked by trypan blue staining (>95% disruption target). The lysate was cleared by centrifugation at 14000g for 1 minute at 4 °C, 2-5 μL was reserved for western blot analysis, and the remainder was aliquoted and flash frozen in liquid nitrogen.

### Antibodies

Primary antibodies used in this study include anti-DDX3 (Bethyl A300-474A; Figure 1), rabbit polyclonal anti-DDX3 (custom made by Genemed Synthesis using peptide ENALGLDQQFAGLDLNSSDNQS; Figure 6; Lee et al. 2008), anti-actin HRP (Santa Cruz Biotechnology, sc-47778), anti-FLAG HRP (Sigma, A8592).

**Figure 1:**
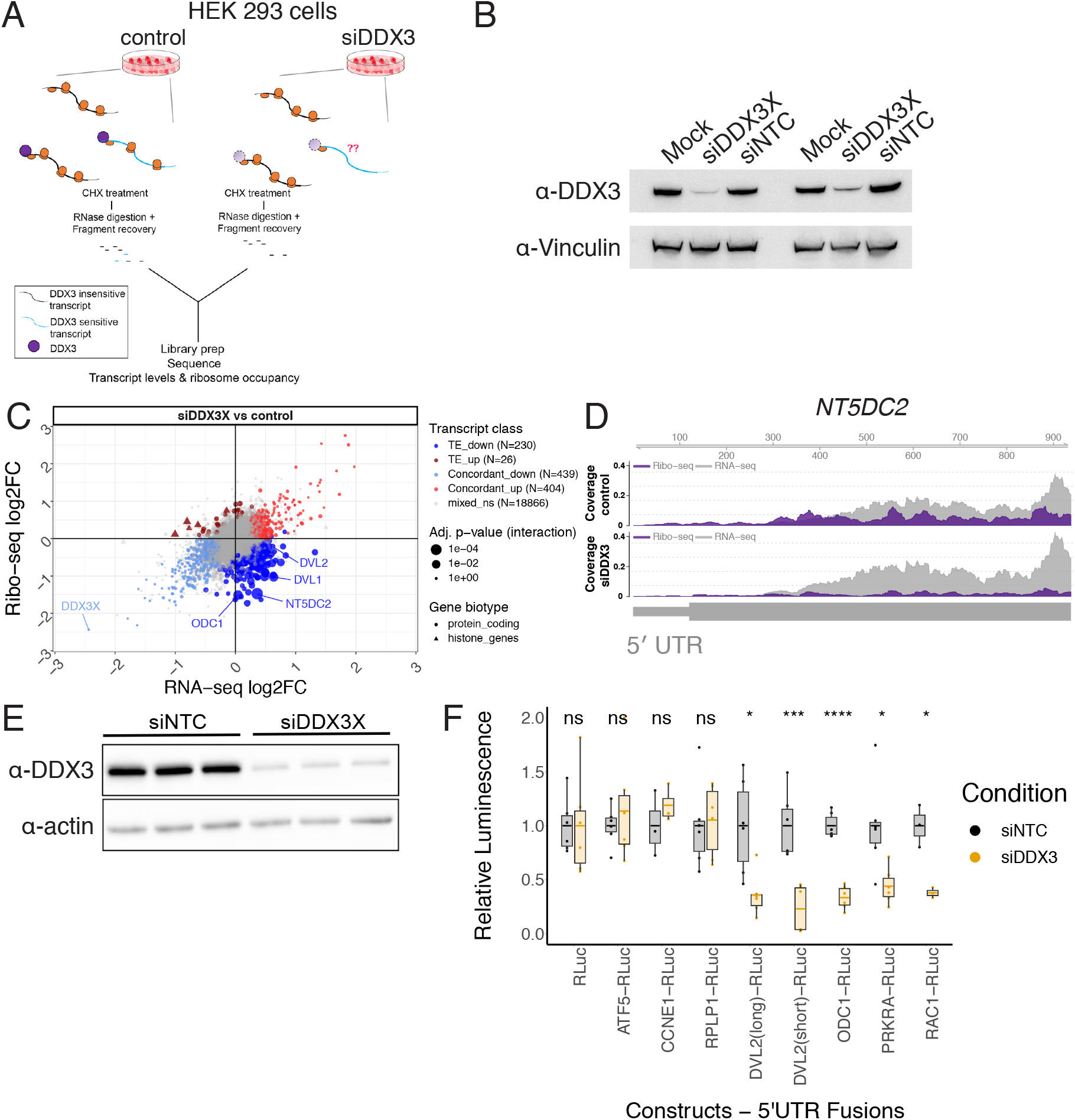
Translational changes upon DDX3 depletion. (A) A workflow of the ribosome profiling experiments. (B) siRNA knockdown efficiency of DDX3 analyzed by western blot. Mock: untreated. NTC: nontargeting control siRNA. (C) The log2 fold change (log2FC) in ribosome profiling or RNA levels are plotted for all genes. Top genes in each category are indicated. Point size indicates the p-value of a significant change in translational efficiency. (D) Tracks showing RNA-seq (RNA) or ribosome profiling (RP) reads that map to the *NT5DC2* gene. *NT5DC2* is an mRNA with differential ribosome occupancy in (C). (E) Western blot analysis of nontargeting (siNTC) or siDDX3 translation lysate samples with antibodies indicated. (F) Renilla luciferase luminescence from in vitro transcribed reporter RNAs translated in vitro in siNTC or siDDX3 HEK 293T lysates.

### *In vitro* translation

5 μL *in vitro* translation reactions were set up with 2.5 μL of lysate and 20 ng total RNA (0.84 mM ATP, 0.21 mM GTP, 21 mM Creatine Phosphate, 0.009 units/mL Creatine phosphokinase, 10 mM HEPES pH 7.5, 2 mM DTT, 2 mM MgOAc, 100 mM KOAc, 0.008 mM amino acids, 0.25 mM spermidine, 5 units RNasin Plus RNase inhibitor (Promega) as described (Lee et al. 2015). Reaction tubes were incubated at 30 °C for 45 minutes, and expression of the reporter was measured using the Renilla Luciferase Assay System (Promega) on a GloMax Explorer plate reader (Promega).

## Results

### Identifying mRNAs that depend on DDX3 for efficient translation

We performed ribosome profiling and RNA-seq to determine the set of transcripts that are affected by depletion of DDX3. *DDX3X* is an essential gene (Chen et al. 2016; Lee et al. 2008), so we transiently knocked down its expression using siRNA and collected ribosome protected footprints (Figure 1A,B). Knockdown efficiencies were ∼90% and ∼70% in replicates (Figure 1B). Measuring changes in both RNA abundance and ribosome occupancy enabled us to distinguish between different modes of DDX3-mediated regulation. We found that depletion of DDX3 affects ribosome occupancy of a minority of messages (Figure 1C). Most changes in ribosome occupancy upon DDX3 depletion were decreases, suggesting that broadly the function of DDX3 is to increase ribosome occupancy (Figure 1C,D). Genes such as *DVL2, NT5DC2*, and *ODC1*, which is described to be translationally-controlled (Steeg et al. 1991), decreased in ribosome occupancy upon DDX3 depletion (Figure 1C, Table S1). Diverse biological pathways were affected by DDX3 depletion, revealing the enrichments of histone mRNAs in the translationally upregulated set, and genes related to neuronal branching in the translationally downregulated set (Figure S1A). We also established a cell line to rapidly and efficiently degrade endogenous DDX3 in human male-derived colorectal cancer HCT116 cells upon addition of an auxin (Figure S1B; Natsume et al. 2016). As with the siRNA knockdown, induced degradation of DDX3 predominantly resulted in decreases in the translation of a subset of cellular messages (Figure S1C). Translation efficiency (TE) changes across the entire transcriptome upon siRNA knockdown and chemical degradation were similar (Figure S1D, Table S1) even though these experiments were performed in different cell lines (HEK293T vs. HCT116) and with different depletion approaches.

Our data suggest that DDX3 directly affects translation of a subset of mRNAs. However, ribosome profiling measures ribosome density, which can be affected by changes to translation initiation, translation elongation, or ribosome stalling. To test whether altered translation initiation contributes to the impact of DDX3 knockdown on ribosome density, we cloned DDX3-sensitive 5′ UTRs from this and previous work (Oh et al. 2016; Chen et al. 2018, 3; Lai et al. 2010) upstream of a *Renilla* luciferase reporter and compared them to a control reporter that is not sensitive to DDX3 depletion. Since *DVL2* has many annotated 5′ UTRs that overlap, we cloned the prevalent isoforms in HEK 293T cells using 5′ RACE, which yielded a short and long isoform (Methods). We then made translation extracts from HEK 293T cells transfected with a nontargeting control siRNA or a DDX3 siRNA (Figure 1E). Next, reporter RNAs were *in vitro* transcribed, capped, and 2′-O methylated and used for *in vitro* translation in wild-type or DDX3-depleted lysate. We found that the 5′ UTRs from the DDX3-sensitive mRNAs *ODC1, PRKRA, RAC1* and *DVL2* also conferred DDX3 dependence to the luciferase reporter (Figure 1F). Therefore, based on these reporter experiments and DDX3 binding pattern, we interpret ribosome occupancy changes upon DDX3 depletion as a result of mis-regulated translation initiation dynamics. However, other reporter RNAs, such as *ATF5, CCNE1*, and *RPLP1*, did not change in the *in vitro* translation upon DDX3 knockdown (Figure 1F). *RPLP1* was identified as an mRNA with uORF occupancy changes upon mutant DDX3 expression in prior work (Oh et al. 2016), while ATF5 and CCNE1 were previously implicated in DDX3-dependent translation (Lai et al. 2010; Chen et al. 2018).

### Defining the features that mediate DDX3-dependent translation

The *in vitro* translation experiments implicated translation initiation and transcript 5′ UTRs in mRNAs that are sensitive to DDX3 depletion. To determine which features contribute to changes in translation upon knockdown of DDX3, we used known translational-control elements to generate a random forest model. A model with 28 features (Methods) was able to moderately predict the translation changes upon DDX3 knockdown (correlation between predicted and observed changes = 0.54, Figure S2A), with few features driving the model performance (Figure 2A, Methods), such as baseline translation levels, GC content in coding sequences and 5′ UTRs, and density of the CERT motif. The CERT motif is a cytosine-rich element that has been implicated in eIF4E- and eIF4A-dependent translation through incompletely understood mechanisms (Jin et al. 2020; Truitt et al. 2015). Interestingly, mRNAs that are sensitive to DDX3 depletion appear to be poorly-translated in HEK 293T cells (Figure 2B). A reduced model only using the four most relevant features performed similarly; conversely, a model built without using sequence predictors or baseline translation levels could not recapitulate translation downregulation effects (Figure S2). 5′ UTR and coding sequence GC content may be indications of increased RNA structure in these regions.

**Figure 2:**
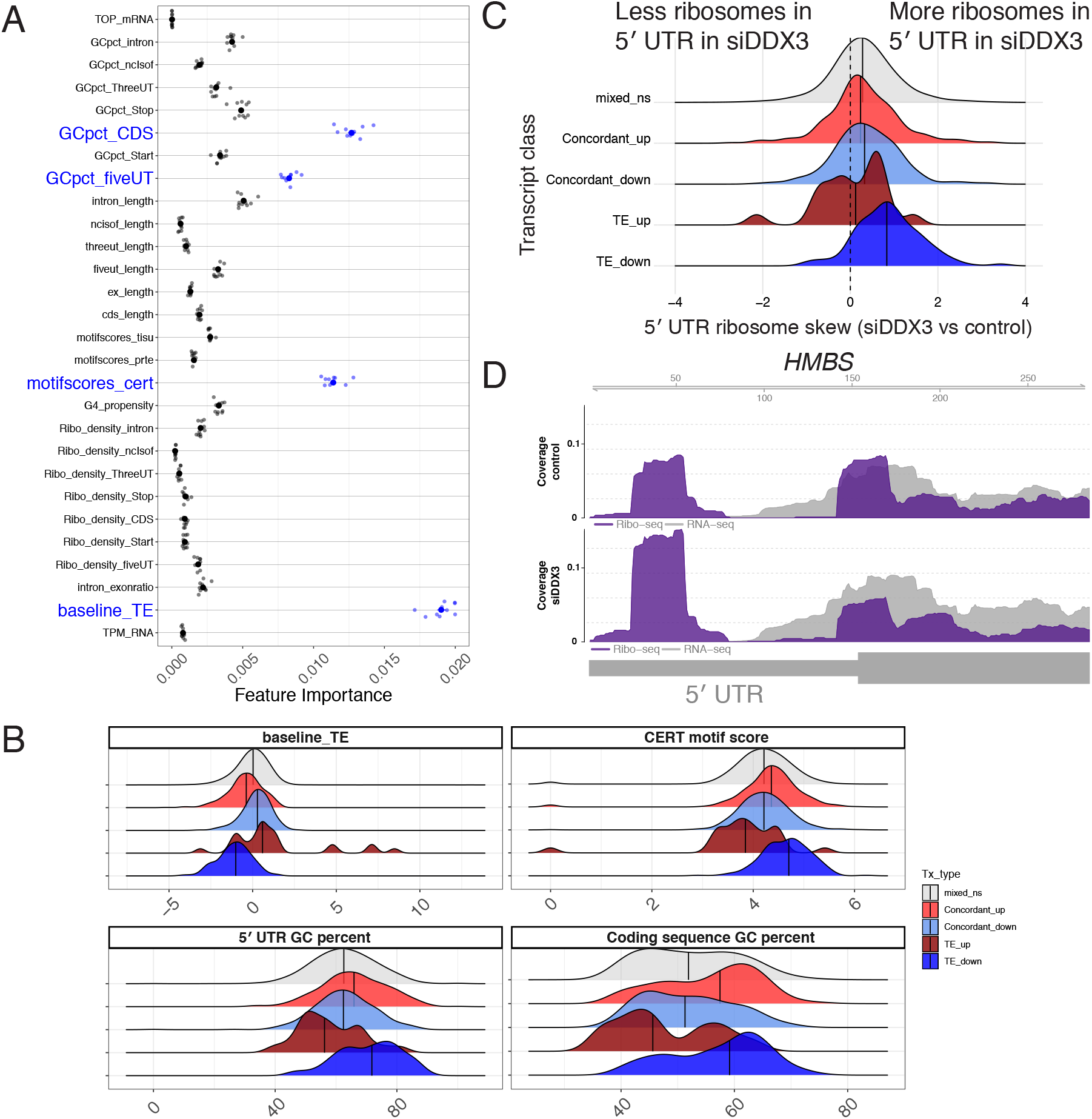
DDX3-sensitive transcripts have complex 5′ leaders. (A) Strength of individual random forest model features that predict TE with R = 0.54 (Figure S2A, Methods). Features in blue are plotted in (B). (B) The translation efficiency (TE) in wild-type cells, CERT motif score, GC-content of 5′ UTRs, and GC content of coding sequences in the indicated gene sets were computed and are plotted as a density plot based on their importance in the random forest model. (C) The fold-change of the ratio in ribosome occupancy in the 5′ UTR versus the coding sequence upon DDX3 depletion as a density plot for transcripts in the indicated sets. A larger 5′ UTR skew value means that there are more ribosomes in the 5′ UTR compared to the coding sequence. (D) An example gene (HMBS) with increases in 5′ UTR ribosomes versus coding sequence ribosomes with tracks as in Figure 1D.

DDX3 is thought to regulate translation through transcript 5′ UTRs, and we found genes that are regulated by their 5′ UTRs such as *ODC1* in the translationally downregulated set (Figure 1E; Auvinen et al. 1992; Steeg et al. 1991). We therefore measured the enrichment of ribosomes in transcript 5′ UTRs, under the hypothesis that depletion of DDX3 might lead to defective scanning and ribosome accumulation (Guenther et al. 2018), or selective ribosome depletion on coding sequences. Indeed, we found more ribosomes in transcript 5′ UTRs relative to coding sequences upon DDX3 depletion (Figure 2C), especially in mRNAs that show translational downregulation. As an example, *HMBS* ribosome occupancy is shown in Figure 2D, which showed changes in ribosome density in its 5′ UTR and therefore may be regulated by upstream ORF (uORF) translation.

### DDX3 crosslinks to ribosomal RNA and 5′ UTRs

The above ribosome profiling and RNA-seq experiments identified the set of transcripts that are affected by DDX3 depletion, but these transcripts could be affected by direct or indirect mechanisms. Therefore, to better define the set of transcripts that are direct targets of DDX3, we measured DDX3 binding sites with high specificity using PAR-CLIP. Previous work using a complementary method (iCLIP) to measure DDX3 binding sites identified 5′ UTR and ribosomal RNA binding. Curiously, even though DDX3 is thought to regulate translation initiation, binding was also identified in coding sequences and 3′ UTRs (Oh et al. 2016; Valentin-Vega et al. 2016). Here, we used the additional specificity afforded by T>C transitions induced by protein adducts on crosslinked uridine residues in PAR-CLIP to examine DDX3 binding across the transcriptome (Hafner et al. 2010).

High-throughput sequencing of RNA fragments crosslinked to DDX3 identified a binding site for DDX3 on the 18S ribosomal RNA (Figure 3; Eliseev et al., 2018). It is possible that these rRNA reads could arise from nonspecific interactions between RNA binding proteins and the highly abundant rRNA. However, while there were many rRNA fragments sequenced following PAR-CLIP, there was only one site spanning nucleotides 527-553 in the 18S with high-confidence T>C transitions (Figure 3B, S3A). This site maps to helix 16 of the 18S rRNA, similar to where Ded1 crosslinks to 18S rRNA in yeast, and does not contain post-transcriptionally modified rRNA nucleotides (Guenther et al. 2018; Taoka et al. 2018). Helix 16 (h16) is on the small ribosomal subunit facing incoming mRNA, which might provide DDX3 access to resolve mRNA secondary structures to facilitate inspection by the scanning 43S complex (Figure 3C). The crosslink site on h16 is just opposite an RRM domain that has been assigned as eIF4B, another factor crucial in ribosome recruitment and scanning (Sen et al. 2016; Walker et al. 2013; Eliseev et al. 2018). This is consistent with observations that eIF4B and Ded1 cooperate in translation initiation on mRNAs (Sen et al. 2016). Recently, it has been proposed that this RRM domain may belong to eIF3g, another translation initiation factor (Brito Querido et al. 2020), which is consistent with a reported interaction between eIF3 and DDX3 (Lee et al. 2008).

**Figure 3:**
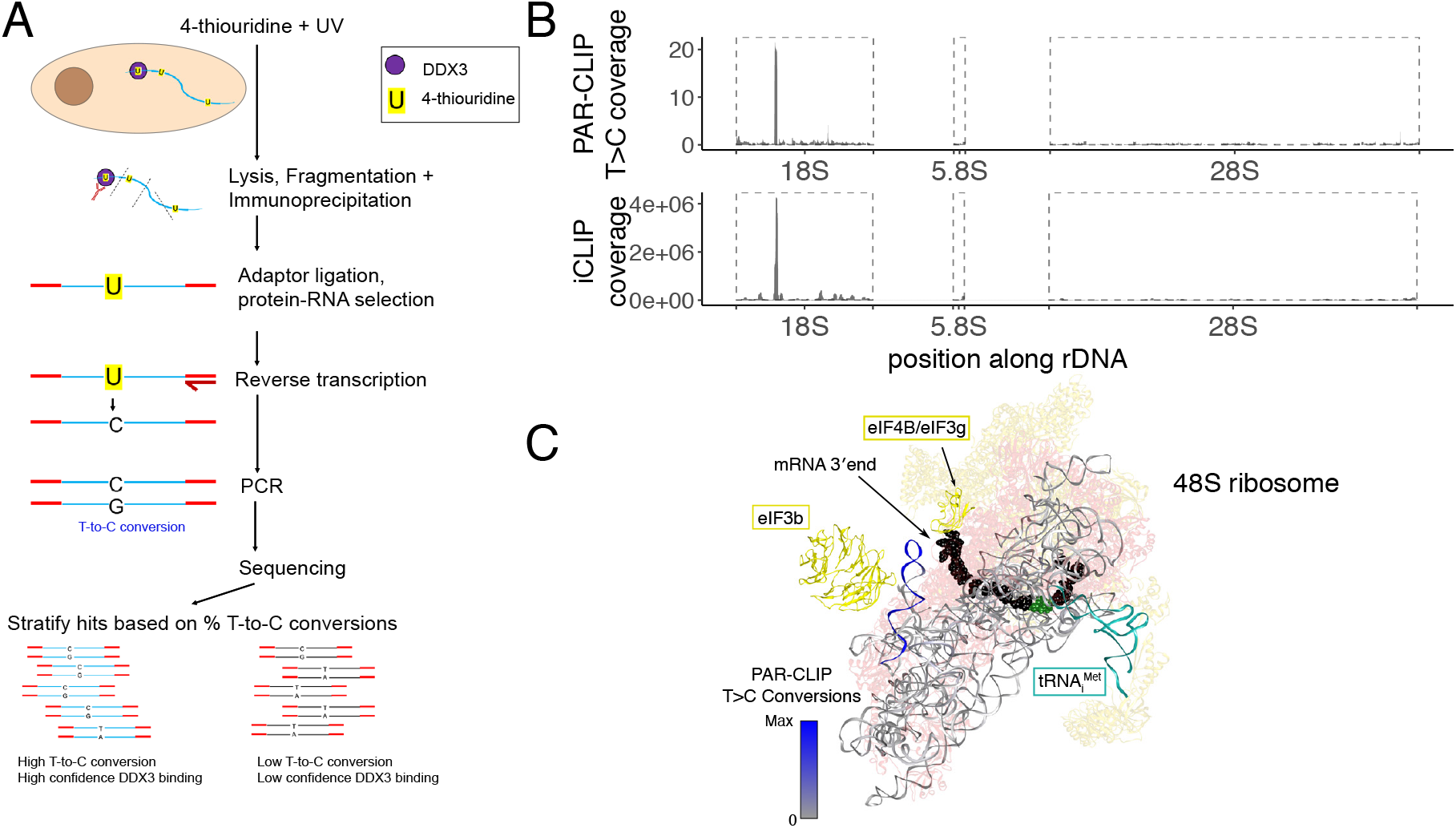
DDX3 binding sites on rRNA identified by PAR-CLIP. (A) A workflow of the PAR-CLIP experiment. (B) PAR-CLIP T>C conversion locations on human rDNA (top) compared to iCLIP coverage from Oh et al 2016 (bottom). Boxed regions refer to processed rRNA transcripts. (C) PAR-CLIP T>C conversion density on the 18S rRNA is visualized from gray to blue on the structure of the 48S ribosome (PDB 6FEC). The peak in (B) is contained in the helix in the upper left in (C), which is h16 of rRNA. Yellow: translation factors; blue-gray: rRNA; pink: ribosomal proteins.

In addition to ribosomal RNA binding, we found that DDX3 interacts primarily with coding transcripts (Figure 4A, Table S2). To identify where DDX3 binds mRNAs, we aggregated peaks across all expressed transcripts in a metagene analysis. We found that DDX3 primarily contacts transcript 5′ UTRs, with a small number of reads mapping in the coding sequence and 3′ UTR (Figure 4B). A large peak was also observed at the start codon, which could reflect kinetic pausing during subunit joining while DDX3 is still bound to the initiating ribosome (Wang et al. 2019). We used available CLIP data to compare the binding pattern of DDX3 to other known mRNA binding proteins (Zhu et al. 2019). We selected three RNA-binding proteins to compare to: eIF3b is a member of the multi-subunit initiation factor eIF3 (Aitken et al. 2016), FMR1 interacts with elongating ribosomes (Chen et al. 2014), and MOV10 is involved in 3′ UTR-mediated mRNA decay (Gregersen et al. 2014; Sievers et al. 2012). The binding pattern of DDX3 most closely resembles the initiation factor eIF3b (Figure 4B). However, we detected some DDX3 binding within coding sequences and even 3′ UTRs, which could arise from background binding, or alternative binding modes. We used the frequency of T>C transitions at each site as a measure of the specificity of protein-RNA interaction (Mukherjee et al. 2018). High specificity crosslinks with frequent T>C transitions resided most often in 5′ UTRs (Figure 4C), as also shown in a translationally-regulated transcript such as *ODC1* (Figure 4D). Confirmation of this binding pattern comes from an independent assay of protein-RNA interaction, as measured by enhanced CLIP (eCLIP) (Figure S3B; Van Nostrand et al. 2016).

Next, we sought to describe the mRNA regions with enriched DDX3 binding. By investigating the sequence-structure context around mRNA peaks in 5′ UTRs, we observed that DDX3 binding sites by PAR-CLIP reside in highly structured regions (Figure 4E). We observed a higher guanine content (accompanied by predicted G-quadruplexes; Figure S3C, S3D) upstream of the binding site; downstream of the peak summit, we detected high GC content resembling the CERT motif (Figure 4F), a regulatory motif highly predictive of DDX3-mediated translation regulation (Figure 2C). Moreover, we observed increased T>C conversion specificity in the 5′ UTR of transcripts whose translation decreases upon DDX3 depletion (Figure 4G), indicative of a possibly stronger protein-RNA association at those regions. Taken together, we conclude that the DDX3 binds and regulates the translation of poorly translated mRNAs exhibiting complex sequence-structure features in their 5’UTRs.

### Identifying translation changes caused by DDX3 mutations

*De novo* genetic variants in *DDX3X* cause developmental delay and intellectual disability in DDX3X-syndrome (Deciphering Developmental Disorders 2015; Snijders Blok et al. 2015; Wang et al. 2018; Lennox et al. 2020). Interestingly, patients carrying inactivating point mutations in DDX3 display more severe clinical symptoms than patients with truncating mutations (Lennox et al. 2020). Point mutations in DDX3 associated with medulloblastoma are dominant negative and act by preventing enzyme closure of DDX3 towards the high-RNA-affinity ATP-bound state (Epling et al. 2015; Floor et al. 2016; Jones et al. 2012; Kool et al. 2014; Pugh et al. 2012; Robinson et al. 2012). This suggests there may be different effects on translation between depletion of DDX3 and inhibition or expression of an inactive mutant.

To test the effect of mutants in DDX3 on translation, we transfected cells with plasmids containing wild-type or mutant DDX3 proteins after auxin-induced degradation of endogenous DDX3, switching expression from the wild-type sequence to an allele of interest (Figure 5A). We used this system to define acute changes to translation caused by DDX3 mutations without allowing the cells to adapt to the presence of inactive DDX3 alleles, which may also be lethal. We measured genome-wide translation changes caused by DDX3 mutants using ribosome profiling (Table S1). A collection of genes related to the double-stranded DNA response were upregulated at the level of RNA abundance, likely due to differences in transfected DNA amounts (Figure 5B, upper right). Broadly, we noticed higher variability in in ribosome occupancy changes (Figure 5B, y-axis) than RNA level changes (Figure 5B, x-axis) when compared to the siDDX3 experiments (Figure 1D), suggesting that one functional difference between mutant and knockdown might involve regulation of RNA steady-state levels. Among the few regulated mRNAs, we observed a robust downregulation of ODC1 translation, which was directly bound by DDX3 (Figure 4D) and strongly downregulated in the DDX3 depletion experiment (Figure 1D).

**Figure 4:**
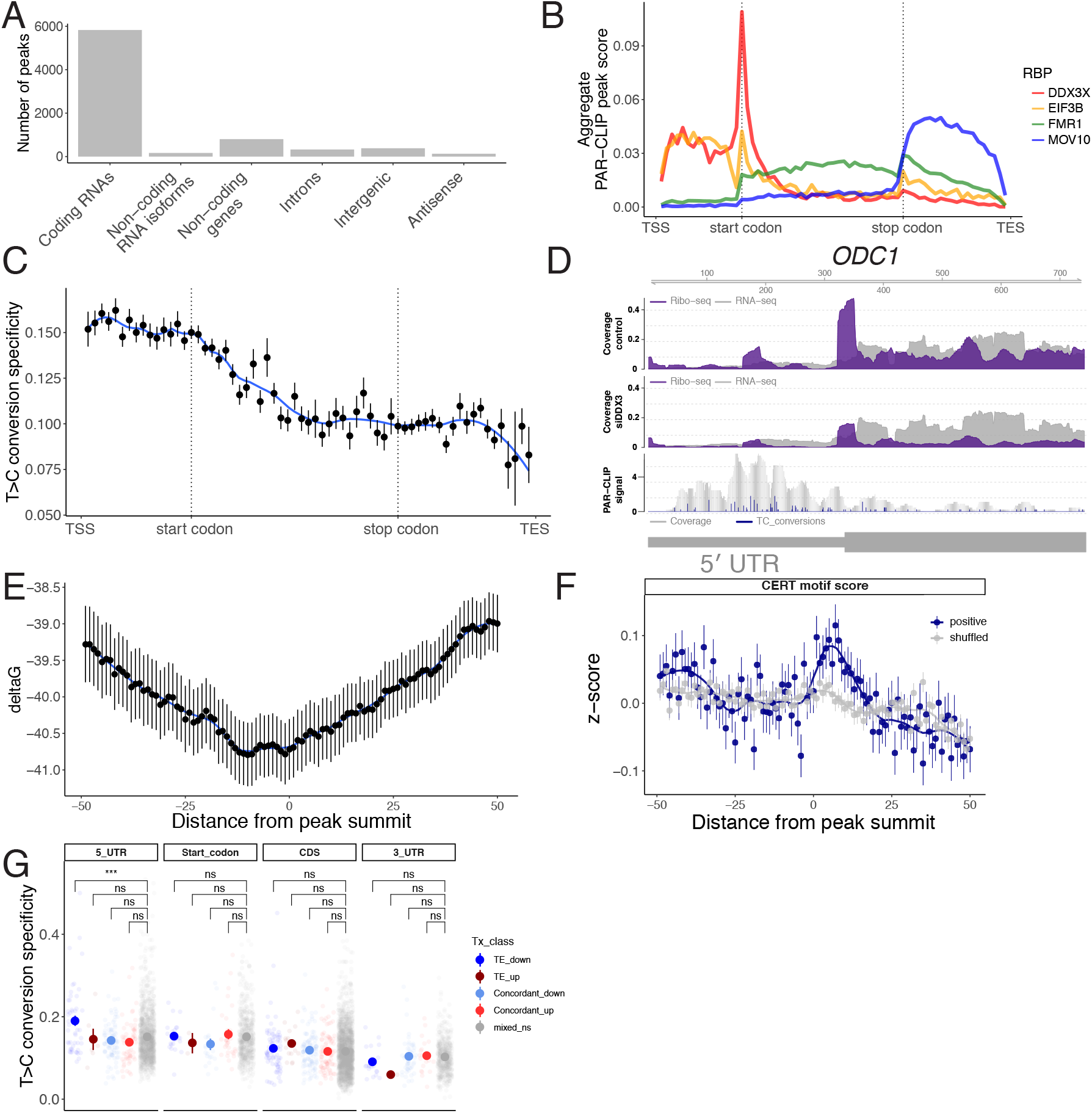
DDX3 binds to structured 5′ leaders of mRNA. (A) Sum of the DDX3 PAR-CLIP peaks mapping on different gene types and regions. (B) A metagene plot of DDX3 PAR-CLIP across all genes and PAR-CLIP data from other RNA binding proteins. eIF3b is a canonical initiation factor, FMR1 binds elongating ribosomes, and MOV10 is a 3′ UTR binding factor. TSS: transcription start site, TES: transcription end site. (C) T>C conversion specificity averaged across all PAR-CLIP peaks across indicated mRNA regions as in (B). (D) PAR-CLIP, ribosome profiling, and RNA-seq across part of the *ODC1* gene. Blue peaks in the PAR-CLIP track indicate T>C conversion events. (E) RNA structure in a window of 100 nucleotides around PAR-CLIP peak summits in 5′ UTRs. (F) CERT motif scores averaged across PAR-CLIP peak summits in 5′ UTRs or for shuffled positions. (G) PAR-CLIP T>C conversion specificity in transcript regions in indicated gene sets from Figure 1.

**Figure 5:**
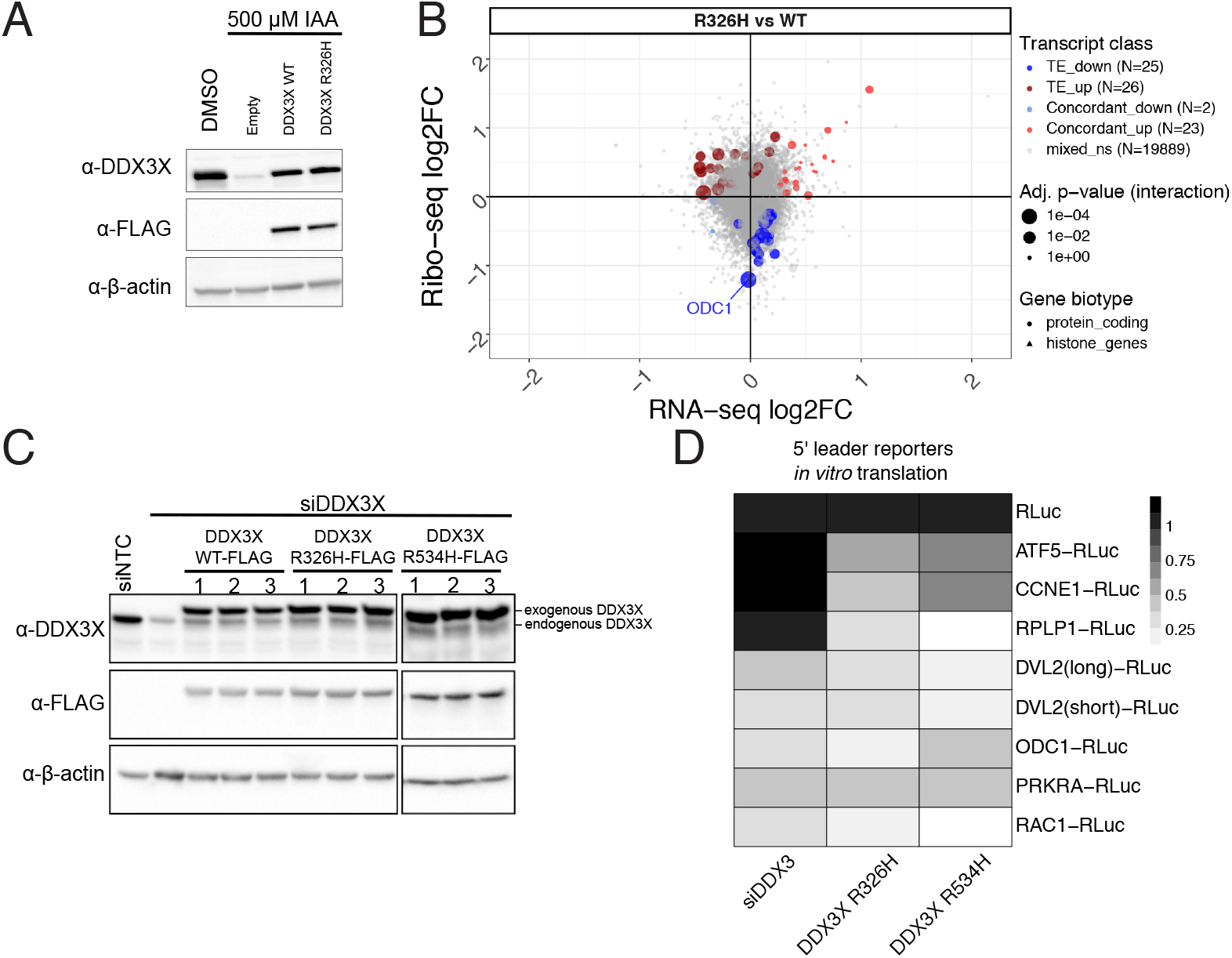
Translation changes caused by R326H mutant DDX3. (A) Western blots of degron DDX3 cells treated with IAA and transfected with empty vector or the indicated constructs. (B) Ribosome profiling and RNA-seq of DDX3 degron cells treated with IAA and transfected with either DDX3 wild-type or R326H mutant. (C) Western blots of cells treated with siDDX3 and transfected with the indicated constructs. (D) In vitro translation performed with indicated reporter RNAs as in Figure 1 in the lysates from panel (C). Reporter sequences are in Table S4.

To further test the difference between knockdown and mutant DDX3 at the level of individual 5′ UTRs, we made *in vitro* translation extracts with wild-type DDX3, R326H DDX3, or R534H DDX3 and measured translation of a panel of reporters (Figure 5C; (Oh et al. 2016). Broadly, we found two classes of reporter RNAs (Figure 5D; S4A, B). One class, including *ODC1, PRKRA, RAC1*, and *DVL2* isoforms decreased in translation in all tested perturbations to DDX3. Another class, including *ATF5, CCNE1*, and *RPLP1* selectively decreased in translation in mutant DDX3 extracts but not in knockdown extracts. We sought to test chemical inhibition of DDX3, as it functionally mimics a dominant negative mutation by blocking ATP binding. We were unable to inhibit translation in a DDX3-dependent manner using RK-33 (Bol et al. 2015), and instead found that it acted as a general translation inhibitor (Figure S4C). Taken together, we found that DDX3 sensitivity for translation is preserved in translation extracts and that depletion of DDX3 appears to have different outcomes on translation than inhibition or dominant negative variants.

## Discussion

*DDX3X* is an essential human gene that is altered in diverse diseases. Here, we use a set of transcriptomics approaches and biochemistry to show that DDX3 regulates a subset of the human transcriptome, likely through resolving RNA structures. We further show that inactivating point mutations in DDX3 yield different outcomes than depletion. Reporter experiments show that 5′ UTRs are sufficient to confer DDX3 sensitivity onto unrelated coding sequences. We conclude that DDX3 affects translation initiation through transcript 5′ UTRs. Our data suggest that the major role of DDX3 is in translation initiation and reveal translation differences between mutated and haploinsufficient DDX3 expression.

We identified binding between DDX3 and helix 16 (h16) on the human 40S ribosome. This is similar to binding sites identified previously using other CLIP approaches (Guenther et al. 2018; Oh et al. 2016; Valentin-Vega et al. 2016), confirmed here using T>C transitions defined by PAR-CLIP. Interestingly, histone mRNAs increased translation upon DDX3 knockdown (Figure 1D), and their translation is dependent on mRNA binding to the rRNA h16 helix (Martin et al. 2016). Our data suggests the possibility that there is competition for h16 between DDX3 and histone mRNAs, and their translation increases upon DDX3 knockdown due to increased accessibility to h16. However, histone mRNAs contain highly repetitive sequences and lack a poly-A tail, perhaps requiring a more tailored approach to precisely investigate the mechanisms of their regulation. The set of mRNAs that require h16 for their translation will be an interesting direction to pursue in the future.

Despite primarily affecting translation, some genes exhibited changes in steady-state RNA abundance upon DDX3 depletion. RNA-level changes could be mediated by indirect effects of a DDX3-dependent translation target or reflect additional mechanisms by which DDX3 regulates gene expression. For instance, DDX3 is an important factor in stress-granule complexes (Shih et al. 2012). Alternatively, RNA-level changes could reflect differences in co-translational RNA decay pathways. Interestingly, we observed more RNA-level changes in DDX3-depletion experiments (Figure 1D and Figure S1B) than in experiments using a mutation in DDX3 (Figure 5B). Potential interactions between DDX3 regulation of both ribosome occupancy and RNA levels will be further explored in future studies.

DDX3 is an abundant protein, with approximately 1.4 million copies per HeLa cell (Kulak et al. 2014), or about half the abundance of ribosomes (Duncan and Hershey 1983). We have interpreted data in this work by hypothesizing that DDX3 is functioning *in cis* by binding to the 40S ribosome and facilitating translation initiation on the associated mRNA. It is also possible that DDX3, alone or in combination with other DEAD-box proteins like eIF4A, functions *in trans* by activating an mRNA prior to 43S complex loading. Future work defining the binding site of DDX3 on the ribosome could enable separation of *cis* and *trans* functions to test these two models, although we note that the functional consequences of DDX3 depletion we have observed here are independent of its functioning in *cis* or *trans*.

DDX3 is altered in numerous human diseases, including cancers and developmental disorders (Sharma and Jankowsky 2014). Some diseases are characterized by missense variants (Jones et al. 2012; Kool et al. 2014; Pugh et al. 2012; Robinson et al. 2012), while others involve predominantly nonsense or frameshift variants (Dufva et al. 2018; Jiang et al. 2015), and still others present with a mixture of variant types (Deciphering Developmental Disorders 2015; Wang et al. 2018; Lennox et al. 2020; Snijders Blok et al. 2015). Our work suggests that variants in DDX3 that deplete protein levels may result in different translation changes than inactivating missense variants. We attempted to directly compare translational changes upon DDX3 depletion identified in this work with previous expression of mutant DDX3 but stopped due to confounding variability in biological sample and library preparation and sequencing protocols. Defining how different mutation types in DDX3 affect gene expression, the underlying molecular mechanisms, and potential therapeutic interventions is an intriguing direction for the future.

## Data access

Sequencing data can be retrieved using GEO accession numbers GSE125114 and GSE157063. Processed datasets and code to reproduce the main figures can be found here: https://github.com/lcalviell/DDX3X_RPCLIP

## Acknowledgments

We thank members of the Floor lab for feedback on the manuscript. Computation was supported by the UCSF Wynton computing infrastructure. This work was supported by the UCSF Program for Breakthrough Biomedical Research, funded in part by the Sandler Foundation (to SNF), the California Tobacco-Related Disease Research Grants Program 27KT-0003 (to SNF) and T30DT1004 (to KW), and the National Institutes of Health DP2GM132932 (to SNF) and F32GM133144 (to SV), and the DFG Priority Project SPP1935 (to KR and ML).

## Author contributions

Conceptualization, L.C., E.W., W.F., M.L., and S.N.F.; Investigation, L.C., K.R., S.V., K.W., and E.W.; Writing – Original Draft, S.N.F.; Writing – Review & Editing, S.N.F., S.V., L.C., E.W., and M.L.; Resources, B.T., M.T., J.K., and W.F.; Funding Acquisition, M.L., and S.N.F.; Supervision, W.F., M.L., and S.N.F.

---

**Figure S1:**
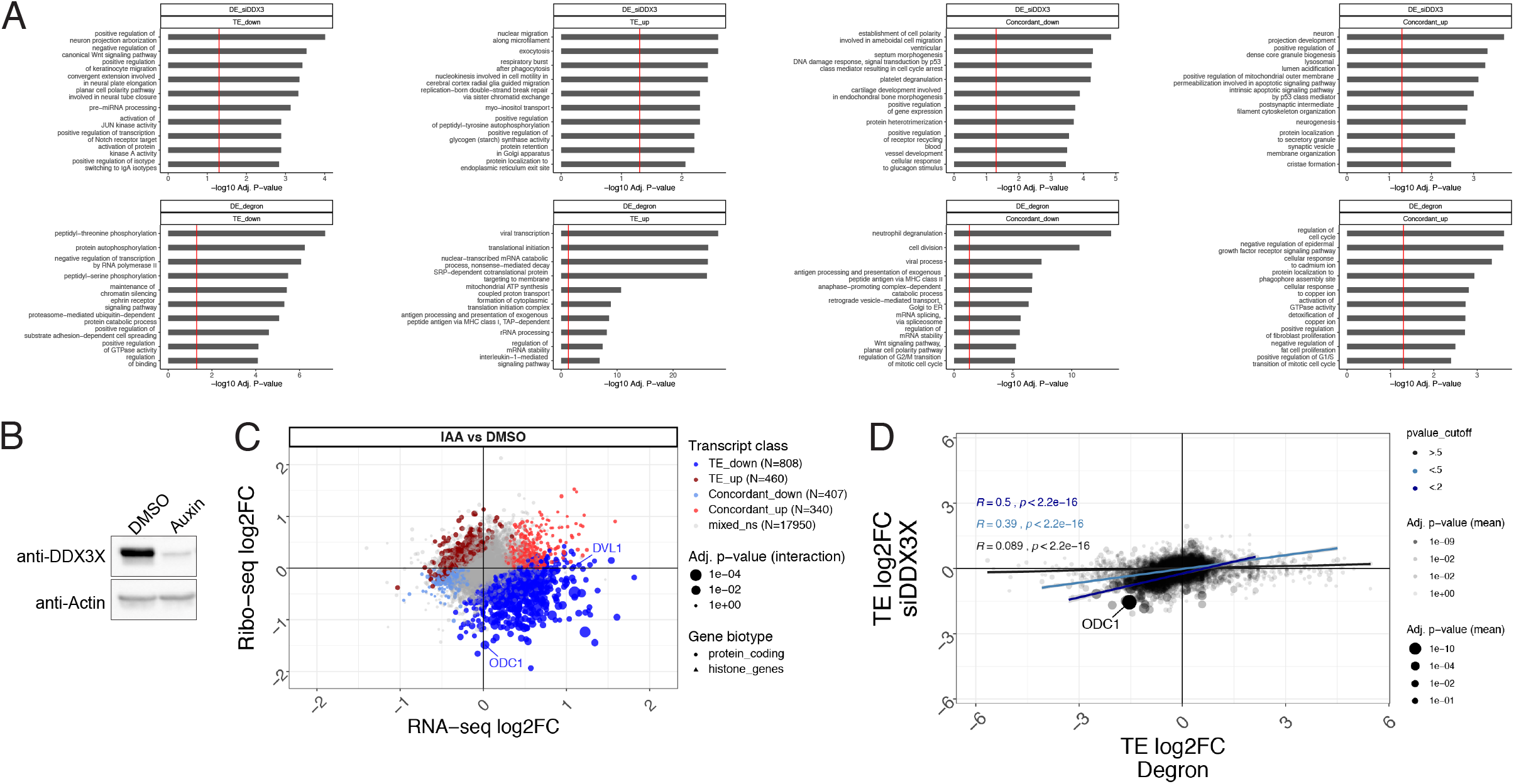
Related to Figure 1 ribosome profiling. (A) Gene ontology analyses for the siDDX3 and degron ribosome profiling experiments. Note that the siDDX3 experiments were in HEK 293T cells while the degron experiments were in HCT 116. (B) Western blot of the DDX3 degron cells treated with IAA (Auxin). (C) Ribosome profiling and RNA-seq fold changes in DDX3 degron (IAA) versus control (DMSO) cells. (D) Translation efficiency (TE) was calculated for all genes in the siDDX3 or DDX3-degron experiments and the two TE values are plotted against each other. ODC1 is indicated. P values for the TE changes are calculated in each contrast using DESeq2 as described in Methods. Correlations are shown at three separate p value cutoffs, where genes with more significant TE changes are more highly correlated between siDDX3 and the degron.

**Figure S2:**
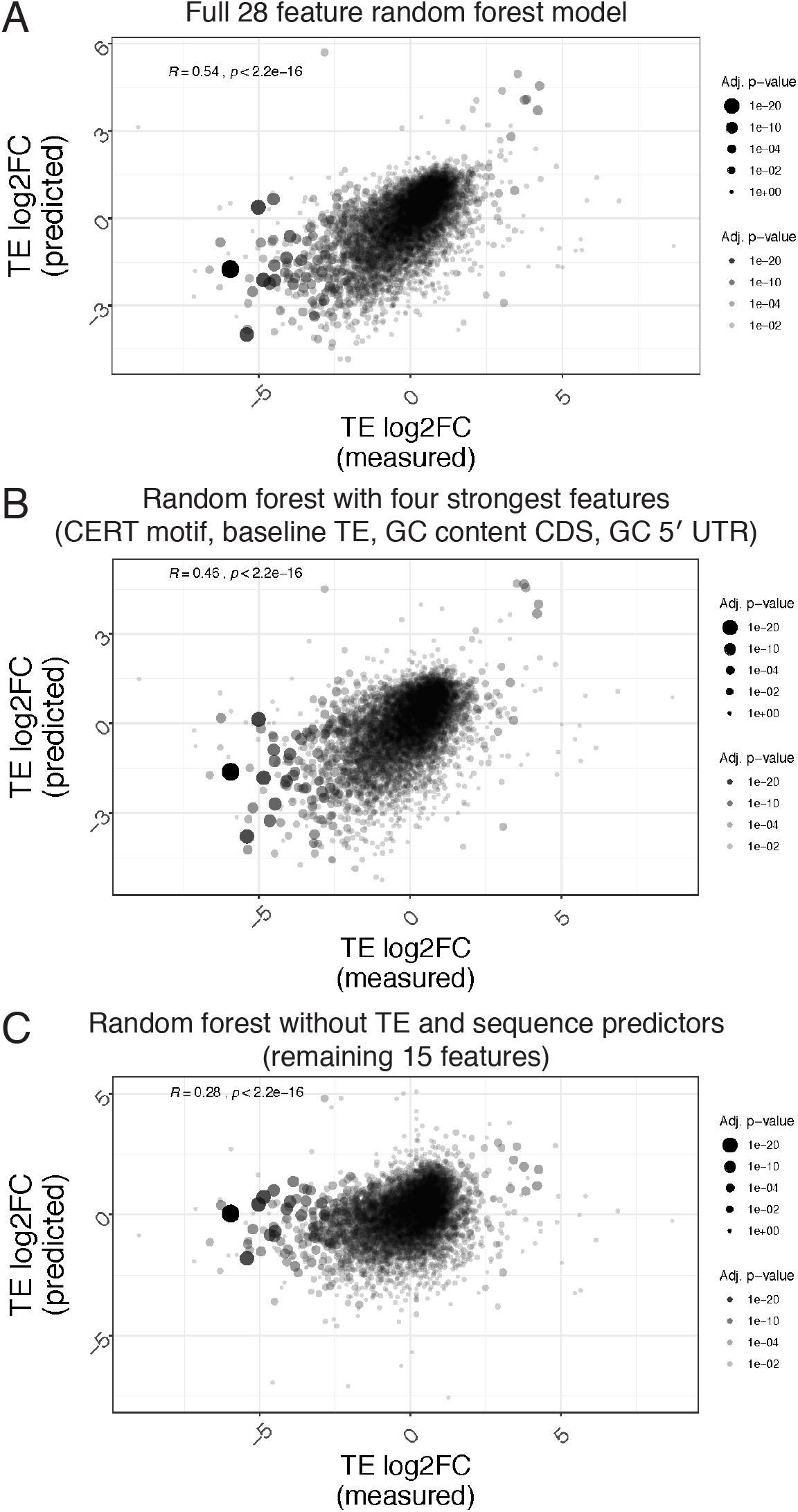
Related to Figure 2 ribosome profiling feature analysis. (A) Correlation between the predicted and observed TE changes, using A) a full Random forest model including all features shown in Figure 2, a B) reduced model with relevant features, or C) less informative features (no baseline TE or sequence features).

**Figure S3:**
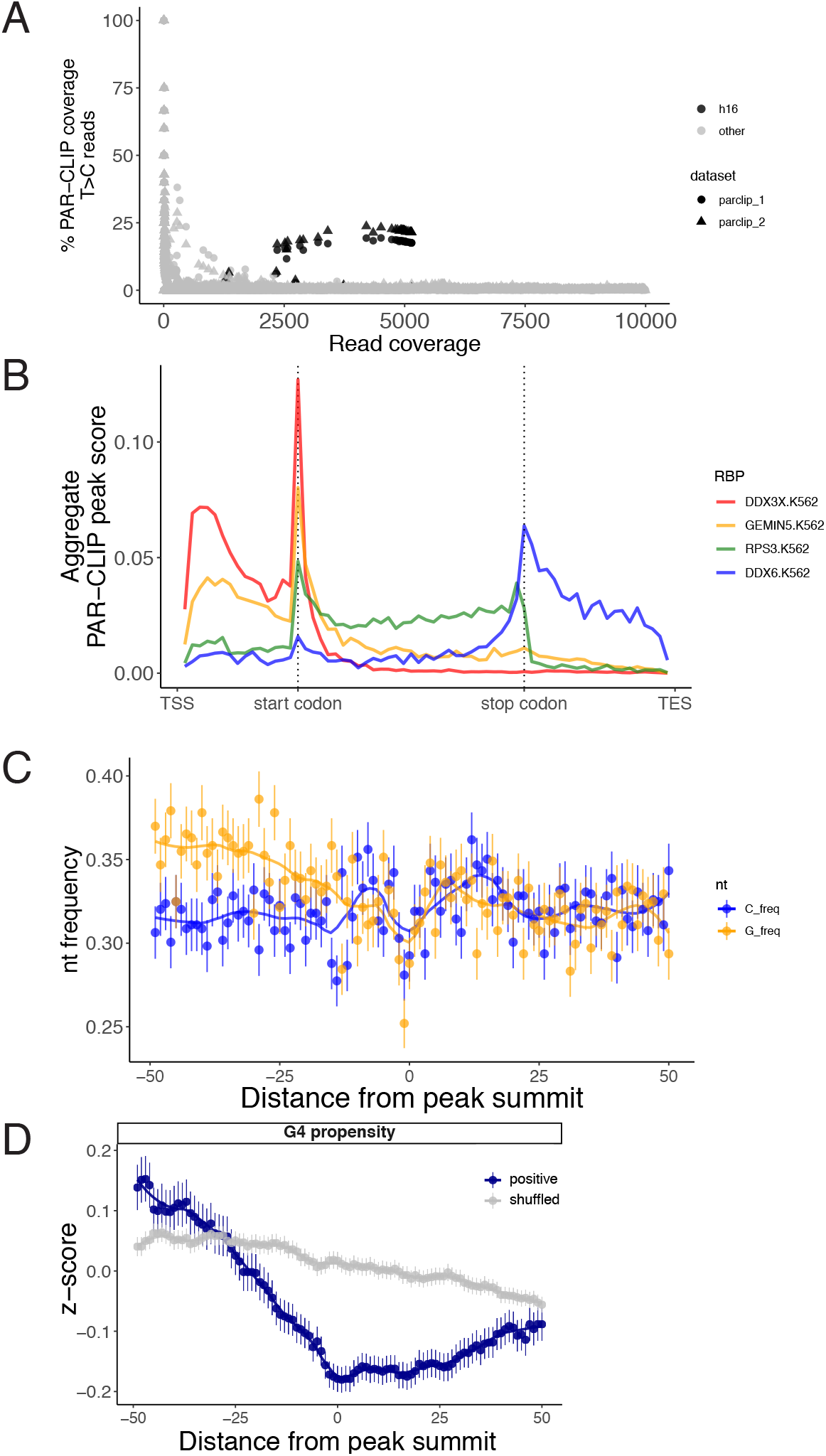
Related to Figure 4 - PAR-CLIP. (A) PAR-CLIP T>C conversion coverage at each rRNA position versus its read coverage for both replicates. Helix 16 (h16) reads have both high T>C coverage and overall read coverage. (B) Metagene analyses for eCLIP datasets from indicated proteins across indicated transcript regions. eCLIP experiments were performed in K562 cells. (C) Guanosine and cytosine frequency averaged across all PAR-CLIP peak summit locations in 5′UTRs. (D) Predicted RNA G-quadruplex propensity of PAR-CLIP sites in 5′UTRs versus shuffled.

**Figure S4:**
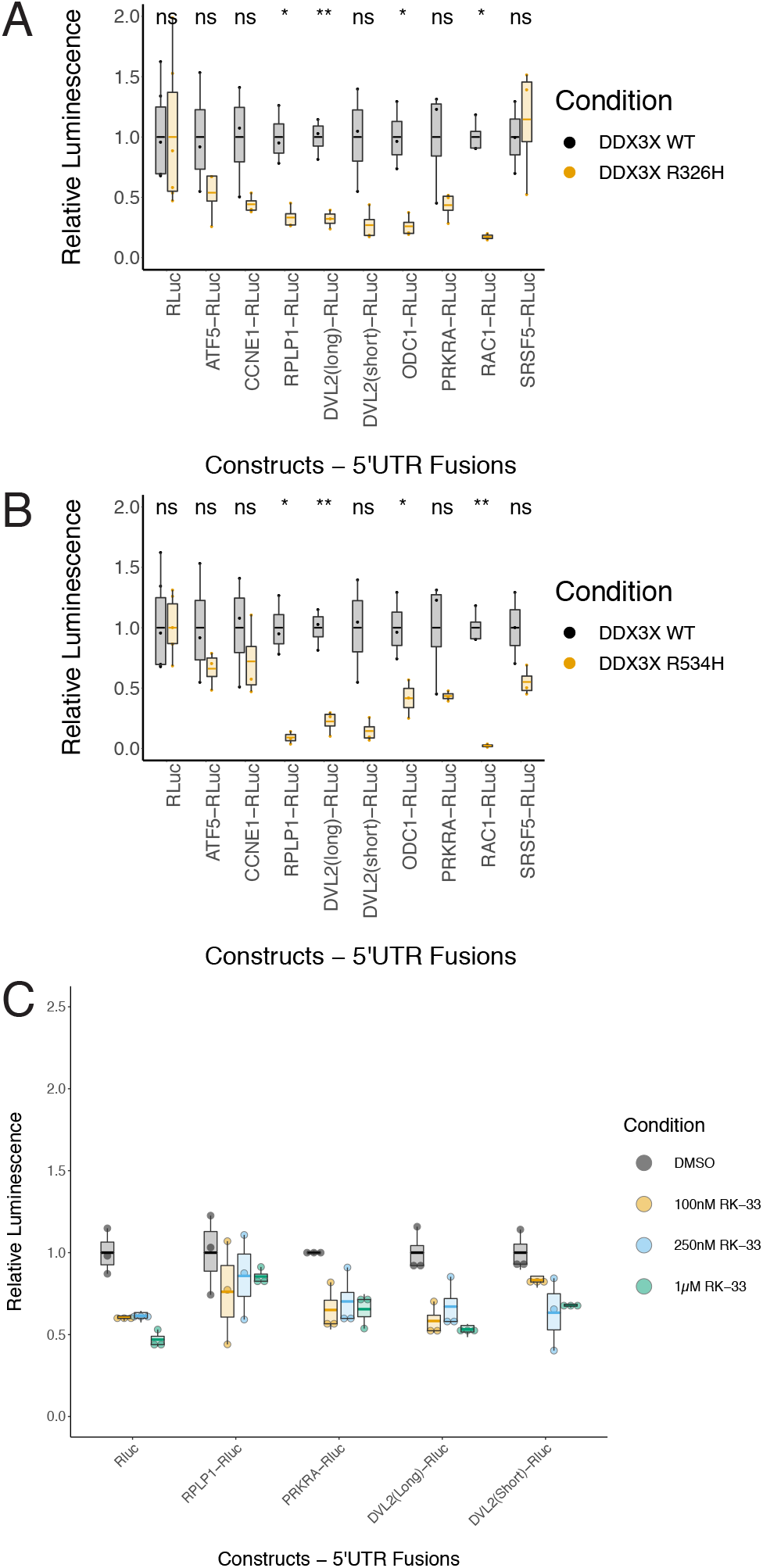
Related to Figure 5 in vitro translation. (A, B) In vitro translation performed with Renilla luciferase fused to the various 5′ leaders indicated. siRNA knockdown of DDX3 complemented with R326H (A) or R534H (B) mutant DDX3X, respectively, compared against siRNA knockdown of DDX3 complemented with wild-type DDX3. (C) In vitro translation with the small molecule RK-33 reveals DDX3-independent inhibition of mRNA translation (compare Rluc control to DDX3-specific 5′ UTR reporters).

