## Supplementary material for "DDX3 depletion represses translation of mRNAs with complex 5′ UTRs": Table S4

Statistics of reporter 5′ UTRs:

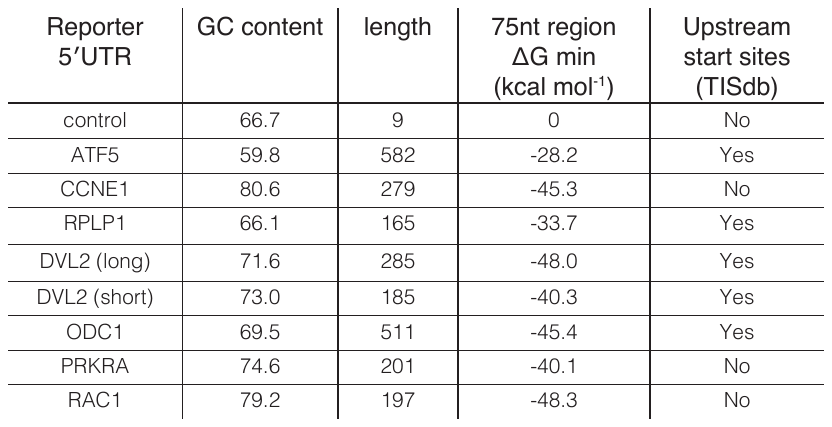

Sequences of reporter 5′ UTRs:

>control

CTAGCCACC

>ATF5_ENST00000595125

ATTTGGGAGAGTGCTGTGACTCATGCTGGACTCTAACCCACGAGGGTTTCTCAGAGTCAGCAGCTGGGGGATGAAGAAGTGAAAAGTGACTGGCAGGAAATTCTGCAAGCAAGGAAAGGGAAAGAGAAATGAACTGGTGCAGGTCTGCGGGAAGAGAATGAGGCTGGATCCTCAAAATCACAGGAGGAAGCAGGCCCAGACCTCAGAGGCAGAAAGAGAAAGAAACCAGAGCTTAGAGTCAGGAGGAGGAAACCAGACCCCGGAGCCACAAGGAGAGGGCTGGATCCCCGGCTCAGAGGGAAGAGTGTCGCCGCCTCTGCCTGCGTAGCCCCGGCCATGGCTCTGTAGCCTCGACCCCTTTGTGCCCCCGGCCCGTCTCCGCGCTCACCACGCCTGCGCTCTCCGCTCCCACCTTCTTTCTTCAGCCGAGGCCGCCGCCGCCTCTCCTTGCTGCAGCCATGGAGTCTTCCACTTTCGCCTTGGTGCCTGTCTTCGCCCACCTGAGCATCCTCCAGAGCCTCGTGCCAGCTGCTGGTGCAGCCTCTCCTGTTGCCATCAGTGCCCAGCACCTGTGCTACAGCC

>RPLP1_ENST00000260379

TGGACACATAAGAGGCTGCGTATAGGCGCGAGAGCCCCTTTCCTCAGCTGCCGCCAAGGTGCTCGGTCCTTCCGAGGAAGCTAAGGCTGCGTTGGGGTGAGGCCCTCACTTCATCCGGCGACTAGCACCGCGTCCGGCAGCGCCAGCCCTACACTCGCCCGCGCC

>DVL2_short_no_accession

TCGCACCCCGCGGCCCGCCCCCCGCCGCCACCCTCGCAGATCCGTGCTTTTTCCCCTTTGCTTCTCTCCCGTACTGGGTCAGTCCTGTCCGCGCTCGCGCGTCGGTTTGCGGGTGTGCGCAGGCGCGGCAGGGGCCATTAGCCCTTTGGGTGGGCGGTGGAGCCCGGGAGCGCGCGGGCGAGACC

>DVL2_long_ENST00000005340

TTAAGTCACGTGACATGAGGAGAGGTGGGCGGGTACCTGGAGGAAGCTCGCGGCGTCGGTGGCGGTGGCGCGCGGCGGCCGCTGAGACCGGGGCTTTGAGTCGCACCCCGCGGCCCGCCCCCCGCCGCCACCCTCGCAGATCCGTGCTTTTTCCCCTTTGCTTCTCTCCCGTACTGGGTCAGTCCTGTCCGCGCTCGCGCGTCGGTTTGCGGGTGTGCGCAGGCGCGGCAGGGGCCATTAGCCCTTTGGGTGGGCGGTGGAGCCCGGGAGCGCGCGGGCGAGACC

>PRKRA_ENST00000325748

GCGAGGGGGCGTAGCCGGAGCTACGGCACCAAGGCTCCGCCCCCACCCTGCCTGCCCCCTCGCTGGAGCAACGCAAGCAGGAGGCGGGGGAGTCGGAGGAGGTGGCGGCGCTGGAGCTCCTCCCGGGGACCAGCGACCCGGGGAGCGAGCACGTCGCTCCGCACCGCTCTTCCTCCAGCCGCTGAGCCGTCCCTTCTCGCC

>RAC1_ENST00000356142

AGTTTTCCTCAGCTTTGGGTGGTGGCCGCTGCCGGGCATCGGCTTCCAGTCCGCGGAGGGCGAGGCGGCGTGGACAGCGGCCCCGGCACCCAGCGCCCCGCCGCCCGCAAGCCGCGCGCCCGTCCGCCGCGCCCCGAGCCCGCCGCTTCCTATCTCAGCGCCCTGCCGCCGCCGCCGCGGCCCAGCGAGCGGCCCTG

>ODC1_ENST00000234111

GACGTCGGCCCGCCGGCGCCCCACCAGCTCCGCGCGGGCCCGGGTTGGCCACCGCCGGGCCCCCGCCCCTCCCCCGGCGGTGTCCCGGCCGGAACCGATCGTGGCTGGTTTGAGCTGGTGCGTCTCCATGGCGACCCGCCGGTGCTATAAGTAGGGAGCGGCGTGCCGTGGGGCTTTGTCAGTCCCTCCTGTAGCCGCCGCCGCCGCCGCCCGCCGCCCCTCTGCCAGCAGCTCCGGCGCCACCTCGGGCCGGCGTCTCCGGCGGGCGGGAGCCAGGCGCTGACGGGCGCGGCGGGGGCGGCCGAGCGCTCCTGCGGCTGCGACTCAGGCTCCGGCGTCTGCGCTTCCCCATGGGGCTGGCCTGCGGCGCCTGGGCGCTCTGAGATTGTCACTGCTGTTCCAAGGGCACACGCAGAGGGATTTGGAATTCCTGGAGAGTTGCCTTTGTGAGAAGCTGGAAATATTTCTTTCAATTCCATCTCTTAGTTTTCCATAGGAACATCAAGAAATC

>CCNE1_ENST00000262643

GAGGGGCTGGGAGCCGCGGCGGGGCGGTGCGAGGGCGGGCCGGGGCCGGTTCCGCGCGCAGGGATTTTAAATGTCCCGCTCTGAGCCGGGCGCAGGAGCAGCCGGCGCGGCCGCCAGCGCGGTGTAGGGGGCAGGCGCGGATCCCGCCACCGCCGCGCGCTCGGCCCGCCGACTCCCGGCGCCGCCGCCGCCACTGCCGTCGCCGCCGCCGCCTGCCGGGACTGGAGCGCGCCGTCCGCCGCGGACAAGACCCTGGCCTCAGGCCGGAGCAGCCCCATC
